# Pulmonary natural killer cells control neutrophil intravascular motility and response to acute inflammation

**DOI:** 10.1101/680611

**Authors:** J. Secklehner, K. De Filippo, J. B. G. Mackey, J. Vuononvirta, X. L. Raffo Iraolagoitia, A. J. McFarlane, M. Neilson, M. B. Headley, M. F. Krummel, N. Guerra, L. M. Carlin

**Affiliations:** Cancer Research UK Beatson Institute, Glasgow, UK; Institute of Cancer Sciences, University of Glasgow, Glasgow, UK; Inflammation, Repair & Development, Imperial College London, London, UK; Life Sciences, Imperial College London, London, UK; Department of Pathology, University of California, San Francisco, CA, USA; Fred Hutchinson Cancer Research Centre, Seattle, WA, USA

## Abstract

The pulmonary immune system defends a huge surface area directly in contact with the contents of the air we breathe. Neutrophils, the most abundant immune cell in the pulmonary vasculature, are critical to immunity but they are also capable of generating life-threatening pathology. Natural Killer cells are the most highly represented lymphocyte subset in the lung, but relatively little is known about their localization, motility or the specific mechanisms by which they contribute to local homeostasis. Here, we used lung-intravital microscopy to directly visualise and quantify neutrophil and natural killer cell dynamics in the pulmonary vasculature of live mice. This approach revealed unexpected sessile behaviour by intravascular natural killer cells. Interactions with natural killer cells made neutrophils scan the endothelium more slowly over larger distances and reduced the number of neutrophils that accumulated in an LPS-triggered inflammatory challenge. This represents a new paradigm by which natural killer cells contribute to lung physiology by diminishing potentially pathogenic neutrophil accumulation.

## Introduction

Gas-exchange in the lungs requires an exceptionally thin barrier between blood and air. To facilitate the diffusion of oxygen and carbon dioxide, the lung has a unique structure that in turn makes it an ideal entrance point for many pathogens (*1*). The immune system must strike a fine balance between sufficient activation and limited tissue damage to provide efficient protection without impairing respiration. Natural Killer (NK) cells are lymphoid-lineage cells of the innate immune system highly-important in the control of viral infections and tumour development (*2, 3*). Interestingly, NK cells are enriched in the lung, making up 10% of pulmonary lymphocytes, compared to 3-4% in the blood, and they are thought to possess a more mature, less cytotoxic phenotype compared to other tissues (*4, 5*). Neutrophils, the most abundant leukocyte in the blood, play a crucial role in initiating the immune defence against many pathogens (*6*–*9*). They are often the first immune cell to be recruited to a site of inflammation and are rapidly activated to kill bacteria and recruit other immune cells by releasing granule contents, extruding DNA ‘traps’ and/or producing inflammatory mediators. This makes neutrophils both capable anti-microbial effector cells and major contributors to tissue damage and pathology (*10*). Until relatively recently, neutrophil regulation has been hard to study, due to their short lifespan and the ease of inadvertently activating them *ex vivo*. In recent years, mainly due to new possibilities for direct cell visualization and advances in genetics, it has been possible to uncover the intricacies of neutrophil behaviour directly *in vivo* (*11*). Crosstalk between neutrophils and NK cells has been suggested before in some conditions in humans and mice (*12*–*15*). Recently, a link between neutrophils and NK cells in cancer has been described whereby neutrophils acquire a pro-tumour phenotype when NK cells are depleted (*16*). Intriguingly, in a model of experimental myocardial infarction, NK cell depletion resulted in increased neutrophilic pathology in the lungs of mice, raising the question of how this influence is mediated (*17*). In fact, the literature on the mechanisms of these putative interactions is somewhat conflicting with indications that NK cells can induce apoptosis of neutrophils or promote their survival (*13, 14*).

A better understanding of the interaction of leukocytes in the lung is required for us to find new pathways to influence the outcome of lung inflammation. Lung pathology in response to infection is mainly caused by neutrophil-driven tissue damage after infiltration but neutrophils are also required for the effective elimination of pathogens and the repair of lung tissue (*18*). Whereas NK cells are important anti-tumour cells (*2*), neutrophils have been implicated in every stage of cancer development including metastasis to the lung (*19, 20*). The potential to interact and the importance of both NK cells and neutrophils in maintaining lung physiology / pathogenesis argued that the behaviour of these cells required closer examination *in situ*. Here, we used lung-intravital microscopy (L-IVM) to directly visualize NK cells and neutrophils in the pulmonary vasculature both in a steady state and during acute lung inflammation to elucidate how crosstalk between these cells is mediated. We performed detailed track interaction analysis on hundreds of individual cell tracks in a series of in vivo imaging experiments to reveal that neutrophil motility differs significantly in cells that interact with NK cells that are sessile in the pulmonary capillaries. The consequence of this interaction is the dampening of the neutrophil response to inflammation in the lung, revealing a previously unappreciated immunoregulatory function for lung intravascular NK cells.

## Results

### Lung NK cells reside within the vasculature where they remain stationary for long periods

NK cells are known to be enriched in the lung compared to other tissues (*3*), however, their precise localisation within the lung has not been well investigated. We used NKp46 (*Ncr1*)^gfp/+^ mice, where NK cells express green fluorescent protein (gfp) (*21*). This mouse allows real-time visualisation of NK cells that are otherwise normal and functional (*22*). We analysed the distribution of NK cells in the tissue versus the vascular compartment in different organs of NKp46^gfp/+^ mice (Fig. 1A-C). To determine the fraction of NK cells that are intravascular we used a differential CD45 staining method (*23*), where fluorescently labelled anti-CD45 is injected intravenously (i.v.) three minutes before humanely killing the mice and then gated on the NK cells by flow cytometry (Fig. S1A). Using peripheral blood as a positive control, we quantified the fraction of NK cells that labelled with i.v. anti-CD45 in the lung, spleen, liver and bone-marrow (BM), tissues where NK cells are reasonably abundant (Fig. 1A-C). Over 90% of NK cells in the lung labelled with i.v. anti-CD45 indicating that they are mostly located in the blood vessels (Fig. 1B and C). As an internal control, we determined that half of the lung T cells did not stain with i.v. anti-CD45, whereas neutrophils showed a similar distribution to NK cells (Fig. S1B-E). Interestingly, although the spleen and BM are highly vascular, the majority of NK cells in these organs were i.v. anti-CD45 negative (Fig. 1C). To confirm intravascular NK cell localization in the lung, we imaged fixed precision cut lung slices (PCLS) (*24, 25*). Confocal microscopy of the slices showed that NK cells were confined within the pulmonary capillaries in a steady state and were not observed in the interstitium, alveoli or perivascular space (Fig. 1D).

**Fig. 1.**
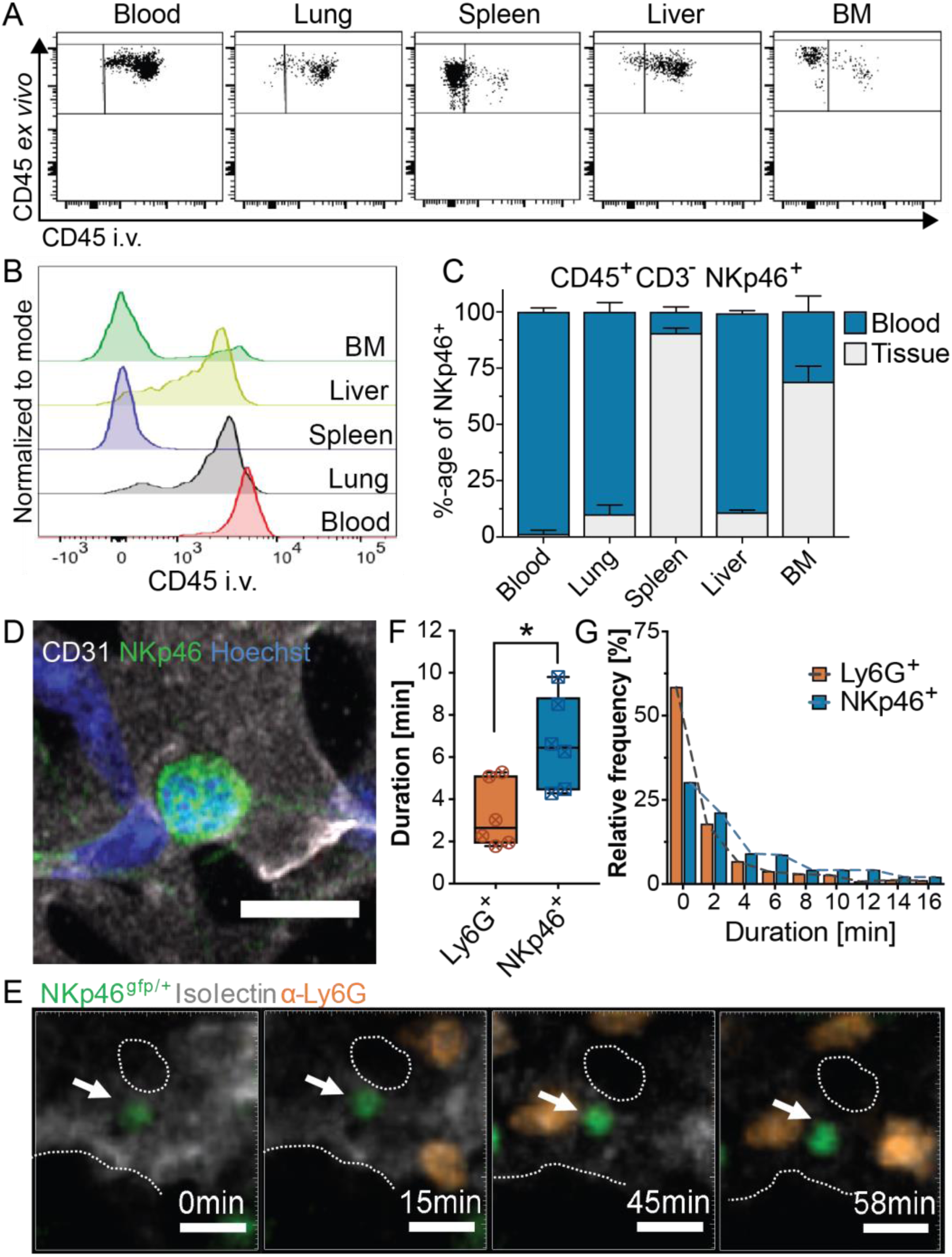
NK cells in the lung are resident in the vasculature. (**A**) Representative plots from flow cytometry analysis of tissues from NKp46^gfp/+^ mice injected i.v. with anti-CD45 Alexa700 and counter stained *ex vivo* with anti-CD45 PerCp Cy5.5. NK cells were defined as CD45, NKp46 positive and CD3 negative. (**B**) Representative histograms of i.v. injected CD45 on NK cells in different tissues. (**C**) Cumulative cytometry data. Error bars represent SD (n=3 mice). (**D**) Representative image of GFP-positive NK cells in the lung vasculature of NKp46^gfp/+^ mice. PCLS were stained with anti-CD31 and Hoechst (Scale bar 10µm). (**E**) Images from L-IVM of unstimulated NKp46^gfp/+^ mice (PBS Ctrl). NKp46-positive NK cells are shown in green (white arrow), anti-Ly6G labelled neutrophils in orange and Isolectin-B_4_ labelled vasculature in grey (dotted lines delineate vasculature; scale bar, 10µm). Comparison of neutrophil and NK cell mean track duration (geometric mean /mouse). (**F**) and frequency distribution of the duration that cells remained in the FOV (**G**) over 20min of L-IVM (n=6 mice; Unpaired t-test with Welch’s correction; \**P* < 0.05).

To obtain kinetic data on the nature of these cells *in situ*, we used L-IVM to study lung NK cells directly *in vivo* (*26, 27*). NK cell tracks were compared to neutrophils that are also found almost exclusively within the lung vasculature in a steady state (*28*). We used small quantities of i.v. injected fluorescently labelled anti-Ly6G, and fluorescently labelled Isolectin (*29*), to label neutrophils and the vasculature respectively. Strikingly, a number of NK cells remained motionless for extended periods of time, with some cells remaining stationary for at least 60 minutes (Fig. 1E; Movie S1). NK cells remained in the field of view (FOV) for significantly longer than neutrophils with a mean NK cell track duration of 6.7±0.9 minutes compared to 3.2±0.6 minutes for neutrophils and over 60% of neutrophils occupied the FOV for less than 2.5 minutes (Fig. 1F and G). These data suggest that the majority of lung NK cells are situated within the blood vessels and individual cells remain within the lung vasculature for long periods. Therefore, we next investigated the potential consequences of this localisation for neutrophils within the pulmonary vasculature.

### NK cells and neutrophils frequently interact in the pulmonary microvasculature

Point-pattern analysis comparing neutrophils and NK cells in fixed PCLS was consistent with both cell types being randomly distributed in the lung (Fig. S2). We hypothesised that interactions between NK cells and neutrophils occurred within the pulmonary capillaries. To test this, we again used L-IVM to image putative NK:neutrophil interactions. Contacts between NK cells and neutrophils spanning several minutes were frequently observed in the pulmonary capillaries (Fig. 2A and C). NK:Neutrophil conjugates were also readily observed in fixed PCLS from naïve mice, fixed before immunofluorescence analysis (Fig. 2B). L-IVM in C57Bl/6J mice showed that neutrophils move erratically through the lung capillaries, frequently stopping to change direction or exiting/entering larger vessels (see Movie S2). In comparison, NK cells were far less motile as established in Fig. 1. Interactions between NK cells and neutrophils occurred frequently, and interactions lasting 10 minutes were not uncommon, while the mean interaction time was 1.85 minutes (Fig. 2C). Quantification of the L-IVM tracks revealed a large number of interacting tracks in each mouse analysed (Fig. 2D). A significantly larger proportion of NK cells interacted with a neutrophil than the proportion of neutrophils that interacted with an NK cell, consistent with the pulmonary NK cells being sessile (Fig. 2E). Interactions were also quantified in PCLS and the percentage of neutrophils interacting with NK cells was comparable to L-IVM result (Fig. 2F). Further analysis of the L-IVM revealed that over 50% of neutrophils interacted with an NK cell more than once (Fig. 2G). Direct cell-cell communication is important for the integrity of the immune system (*30*). To determine whether interactions with NK cells could have functional consequences for the neutrophils we performed more detailed analysis of the interactions.

**Fig. 2.**
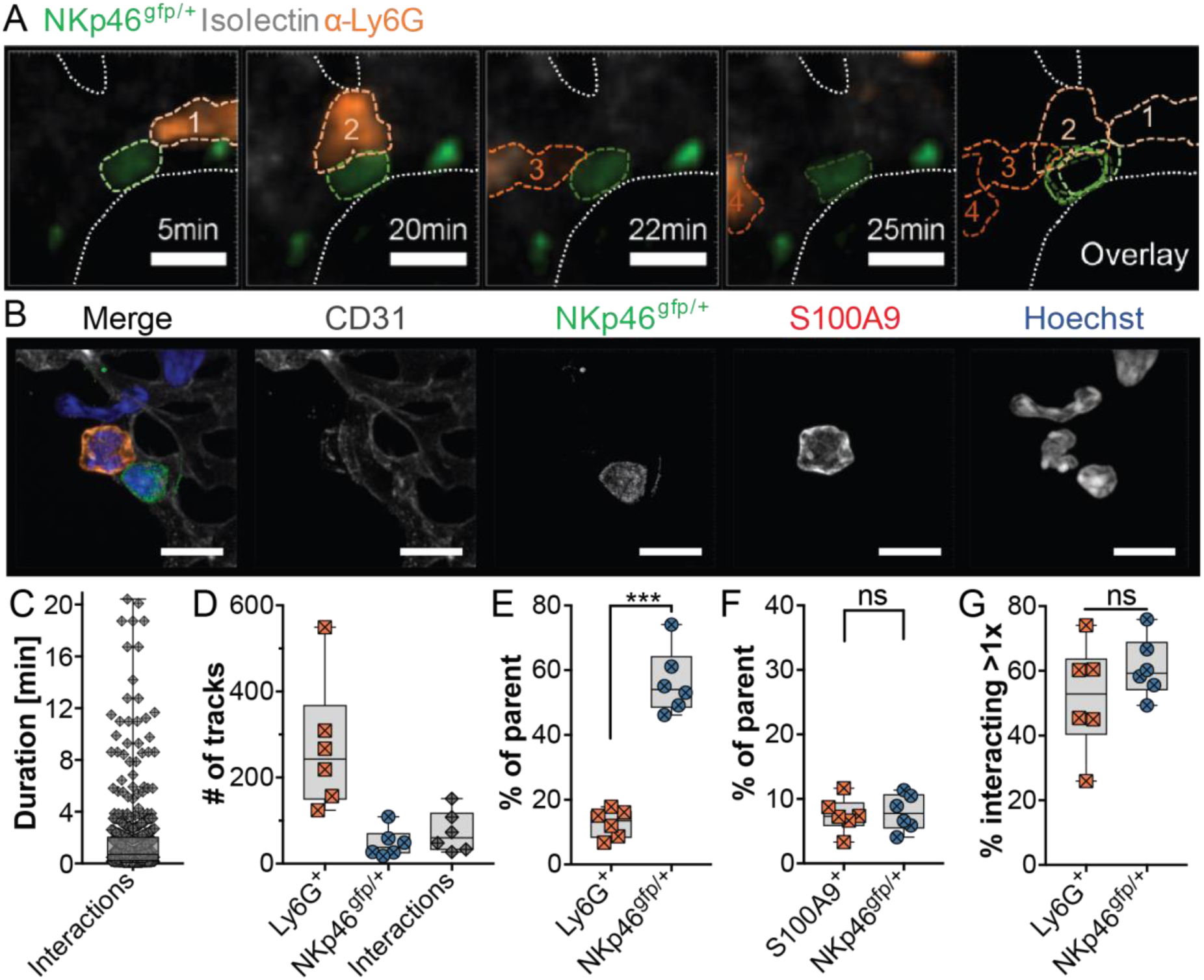
NK cells and neutrophils form long-term interactions in the pulmonary capillaries. (**A**) L-IVM of NKp46^gfp/+^ mice injected with anti-Ly6G to label neutrophils (orange). Dotted lines images represent cell movement over time. (scale bar 10µm; gaussian filter 1.01µm). (**B**) PCLS slices were stained with anti-CD31 and anti-S100A9. Images of S100A9 positive neutrophil:NKp46-GFP positive NK cell conjugates in PCLS (scale bar 10µm; gaussian filter 0.07µm). (**C**) Duration of individual interactions lasting over 3.4 seconds (Mean=1.85; n=6). (**D**) Track numbers from 20 min of baseline L-IVM imaging, excluding tracks shorter than 3.4 seconds (n=6 mice). Percentage of neutrophils or NK cells interacting during baseline imaging (**E**) or in fixed PCLS (**F**) (n=6 mice, paired Student’s t-test). (**G**) Fraction of total neutrophils or NK cells interacting more than once during 20min of L-IVM (n=6 mice, paired Student’s t-test, *** *P* < 0.0005).

### Cellular material is transferred from neutrophils to NK cells during cell:cell contact in the pulmonary capillaries

We inferred that ‘information’ might be exchanged between NK cells and neutrophils in pulmonary capillaries during their cell:cell interaction. Close examination of our L-IVM data supported this notion, by revealing occasional ‘patches’ of anti-Ly6G Ab labelled material being transferred to NK cells during contact with a neutrophil, consistent with the transfer of a membrane domain or vesicle rather than ‘membrane-mixing’ (Fig. 3A; Movie S3). To obtain higher resolution images of the transfer, we performed imaging in live PCLS (Fig. 3B) measuring the average size of transferred fractions at 1.15±0.27μm by 0.84±0.16μm (Fig. 3C). It is unclear how Ly6G could contribute directly as Ly6G-deficient mice exhibit no phenotype in response to LPS (*31*). We hypothesised that Ly6G could just be a GPI-linked (*32*) ‘by-stander’ enabling the visualisation of the transfer, while other material is also exchanged. To this end we performed *in vitro* imaging of isolated neutrophils labelled with a lipid membrane dye, co-cultured for up to 3 hrs with lung NK cells, revealing NK cells with clear dye positive patches of the label originally on the neutrophils only (Fig. 3D). Taken together these data suggest a coregulatory activity mediated by membrane capture by the NK cells perhaps analogous to the ‘trogocytic’ regulation exhibited by other lymphocytes (*33*–*35*).

**Fig. 3.**
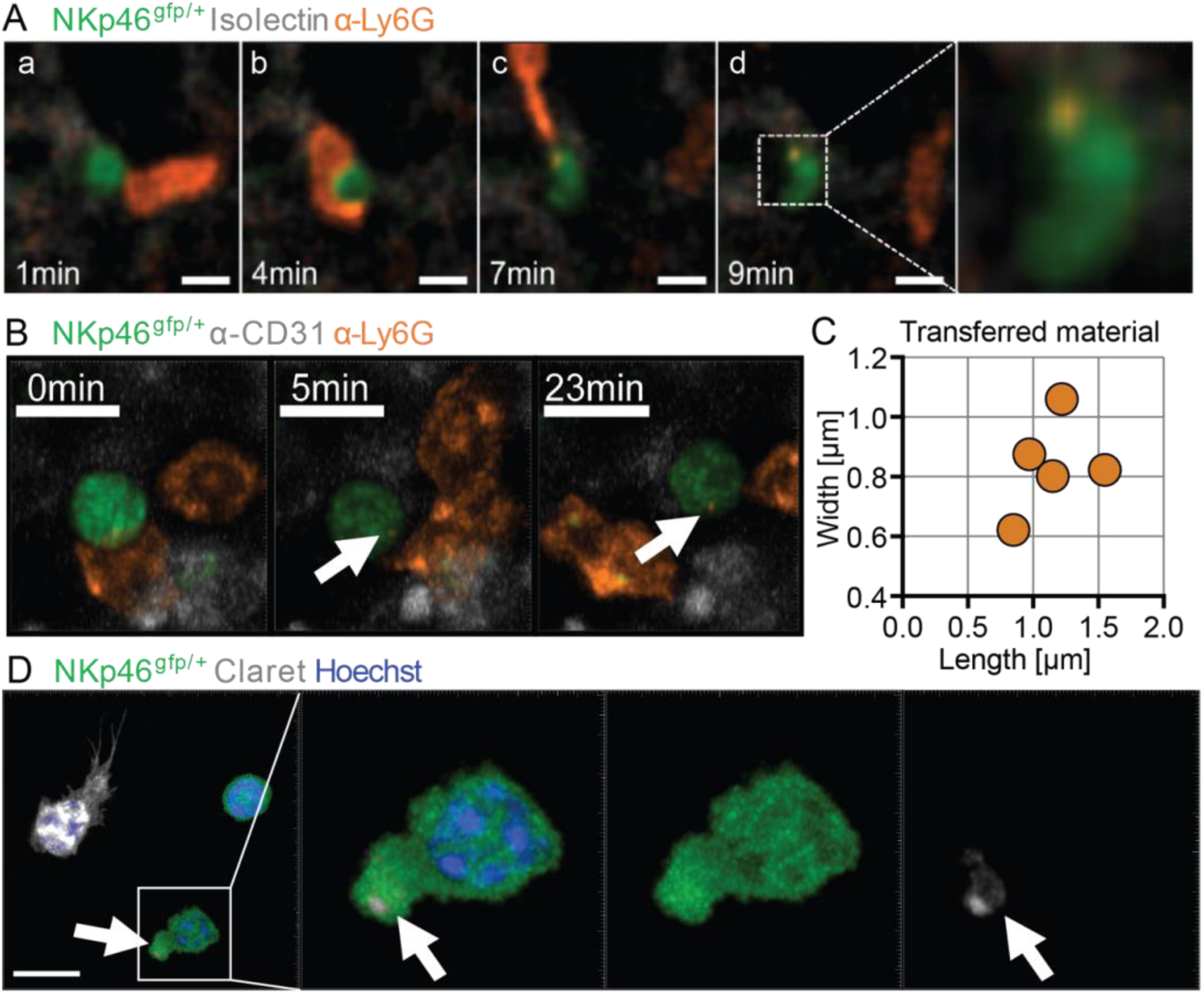
Cellular material is transferred from neutrophil to NK cells. (**A**) Images of L-IVM time-lapse sequence (a-d) from a NKp46^gfp/+^ mouse injected with anti-Ly6G and Isolectin-B_4_. Close contact between an NK cell and a neutrophil in the vasculature with subsequent intercellular transfer (scale bar 10µm; gaussian filter 1.01µm). (**B**) Confocal imaging of live PCLS from NKp46^gfp/+^ mouse. Cells were labelled with anti-Ly6G and anti-CD31 (scale bar: 10µm) (**C**) Area of transferred material in live lung slices (n=5 mice from independent experiments). (**D**) Live imaging of neutrophil and NK cell co-culture. NK cells (green) were isolated from the lungs of NKp46^gfp/+^ mice. Neutrophils were isolated from the blood of the same mouse and labelled with Claret CellVue cell surface dye (grey) and Hoechst to label nuclei (blue). Representative images are shown (scale bar = 10 µm), arrows point to transferred material.

### Interactions with NK cells change neutrophil behaviour

To determine the effect of these direct interactions we performed detailed and extensive track analysis across all mice imaged (Fig. 4) to compare neutrophils that interacted with NK cells with those that did not, in each mouse. Strikingly, neutrophils that were not observed to interact with an NK cell had significantly shorter tracks in terms of both duration and length, increased speed, and displaced less far from their origin (Fig. 4A, B; Table S1). Importantly, when these tracks where further dissected into ‘before’, ‘during’ and ‘after’ interaction sequences (Fig. 4C; Table S2), the interacting neutrophil parameters were not significantly changed, although there was a trend towards reduced displacement during the interaction since the NK cells remain sessile (Table S2). Therefore, neutrophils that interacted with NK cells have altered motility compared to those that do not and that this cannot be accounted for just by differences in the ‘during’ phase alone. In summary, the neutrophils that interacted with NK cells moved more slowly and could be tracked over longer distances, effectively scanning the endothelium more slowly and this could not be accounted for simply by them slowing down as they interact with NK cells.

**Fig. 4.**
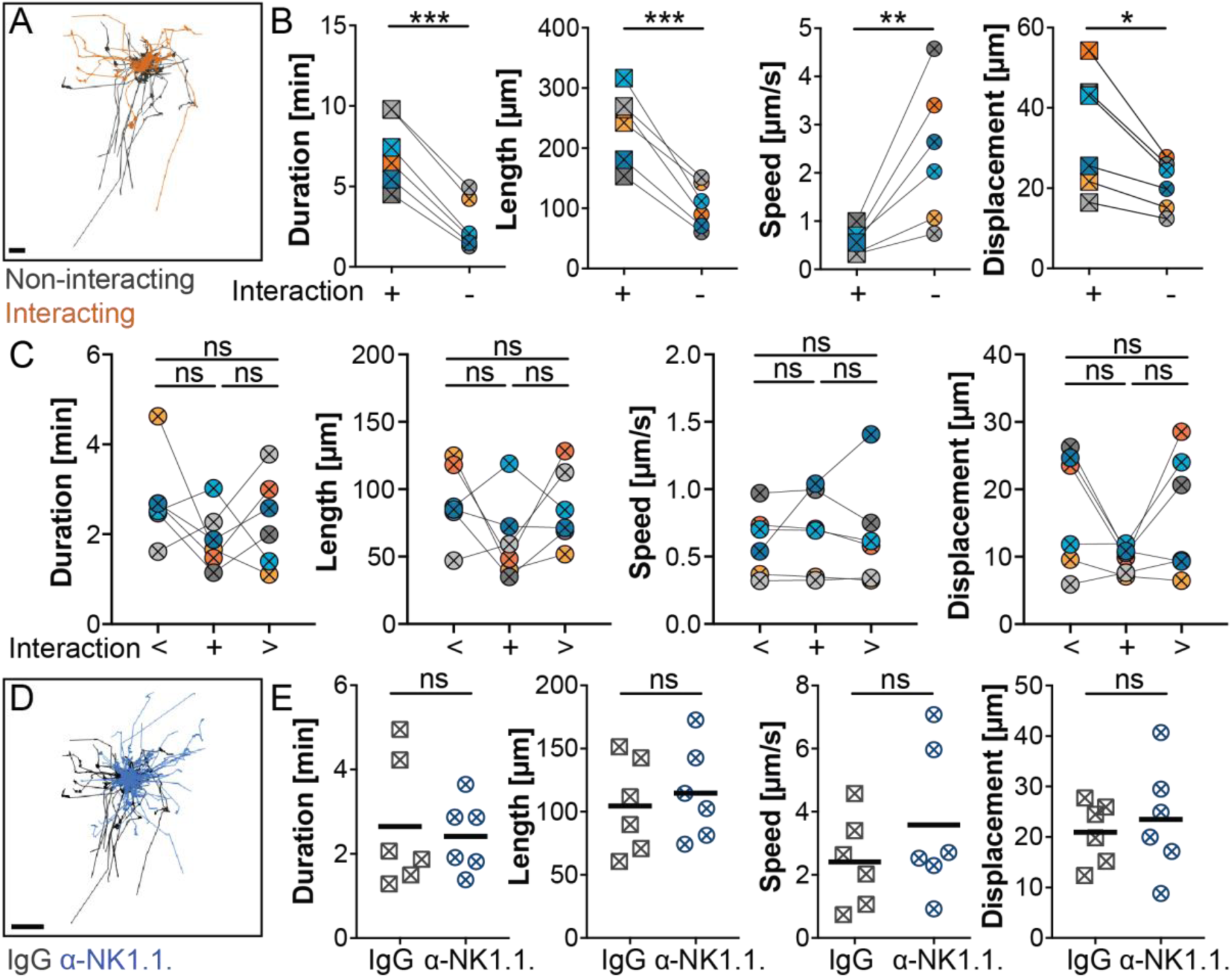
NK-interacting neutrophils exhibit slower motility tracked over longer distances, but NK cell depletion does not alter bulk steady state neutrophil kinetics. (**A**) Representative image of cell tracks of neutrophils that do and don’t interact with NK cells, given a common origin (scale bar: 50µm). (**B**) Track duration, length, speed and displacement compared in NK-interacting versus non-interacting neutrophils in the same mouse. Arithmetic means from each mouse are displayed. (n=6 mice; paired t-test; \**P* < 0.05, \*\**P* < 0.005, \*\*\**P* < 0.0005). (**C**) Interaction analysis showing neutrophil track duration, length, speed and displacement before (<) during (+) and post (>) interaction with an NK cell (n=6 mice; data points represent arithmetic means of tracks; paired Student’s t-tests). (**D**) Representative image of cell tracks from IgG Control and NK cell-depleted mice, given a common origin (scale bar: 50µm). (**E**) Neutrophil track duration, length, speed and displacement were compared between NK cell-depleted and IgG Ctrl mice (n=6 mice, line represents mean; unpaired t-test with Welch’s correction).

To further understand what role NK cells could play in the regulation of neutrophils we depleted NK cells using an established Ab-depletion protocol (i.p. injection of anti-NK1.1.; PK136 (*36*); Fig. S3A). After depletion, flow cytometry confirmed that NK cells were virtually absent in the blood, lung and spleen (Fig. S3B). However when we used this ‘bulk approach’ we couldn’t observe significant differences in the overall number of neutrophil tracks or track parameters (Fig. 4D, E; Table S3) in NK cell-depleted compared IgG treated mice and levels of CXCL1 and CXCL2 were comparable (Fig. S3C and D) as was CXCR2 expression on neutrophils (Fig. S3E and F). These results show that NK cell depletion itself did not cause low level inflammation which could alter neutrophil responses, an important control for the subsequent analysis of neutrophil behaviour during endotoxin induced lung inflammation.

### NK: Neutrophil interactions diminish the early neutrophil response to inflammation

To better understand the role of these interactions, we studied neutrophil behaviour in an established model of acute lung inflammation by airway endotoxin (lipopolysaccharide; LPS) application (*37*). Neutrophils are known to respond rapidly to LPS application, they accumulate in the lung vasculature and show increased surface expression of CD11b within 30 minutes (*28, 38*). L-IVM was performed in NK cell-depleted NKp46^gfp/+^ mice or IgG controls treated with LPS or PBS intratracheally during imaging (Fig. 5A and B). LPS instillation led to a rapid increase in the number of neutrophils as expected, but approximately 50% more neutrophils accumulated in the pulmonary capillaries imaged in NK cell-depleted mice at this very early timepoint (Fig. 5B, Movie S6). Although the gradient of the initial increase in our L-IVM experiments showed that neutrophils were not arriving more rapidly, the neutrophil number plateaued at a greater fold-change from baseline suggesting accumulation via reduced rate of neutrophil egress rather than increased infiltration per se.

**Fig. 5.**
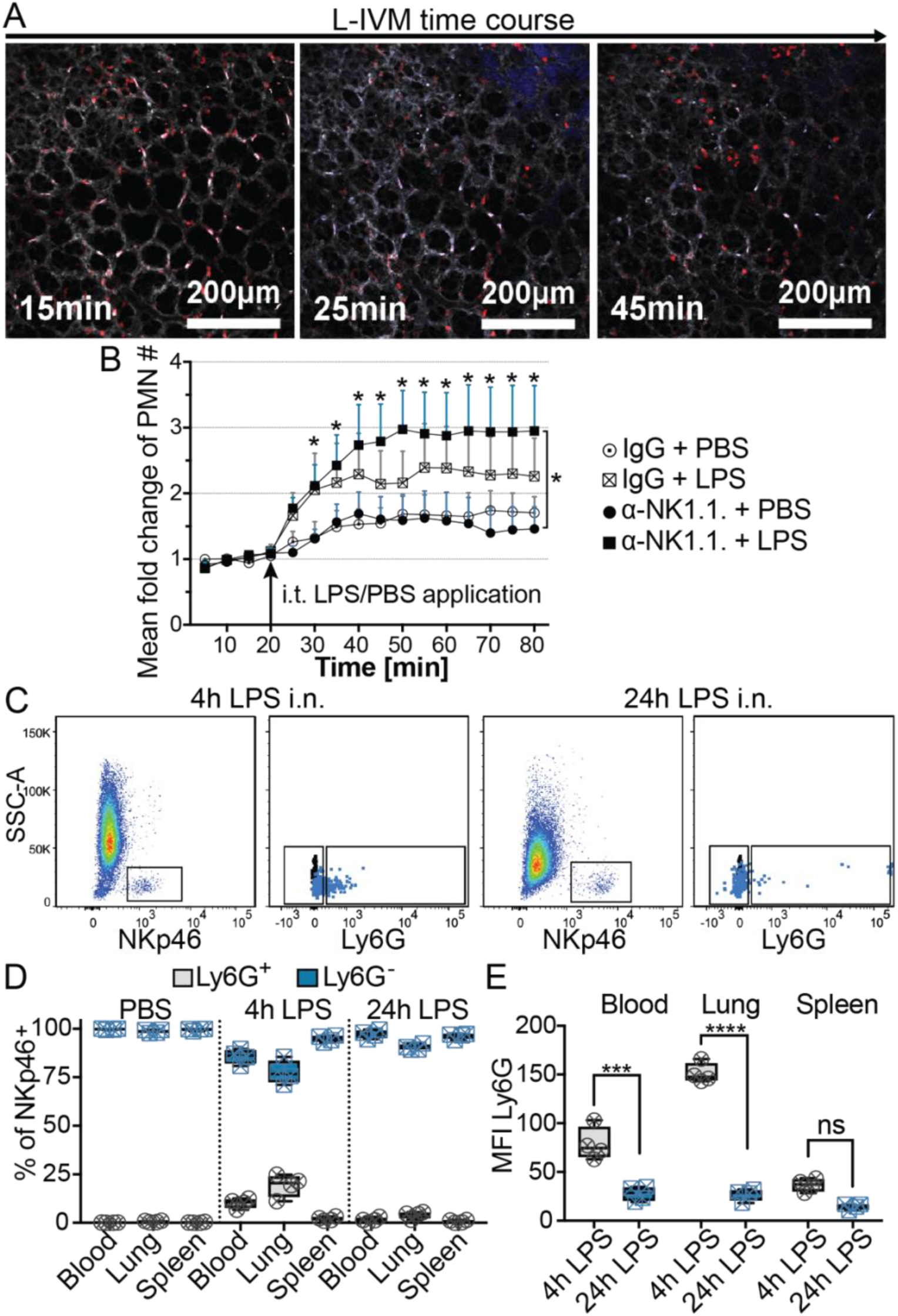
NK cell depletion results in increased neutrophil accumulation in response to LPS-induced lung inflammation. (**A**) Representative images of L-IVM in a C57Bl/6J mouse. The vasculature and neutrophils were labelled by i.v. injection of Isolectin-B_4_ (grey) and anti-Ly6G mAb (red). Acute lung inflammation was induced by intratracheal application of LPS in PBS supplemented with Isolectin-B_4_ (blue) after 20 minutes of imaging to establish a baseline (n=3 mice). (**B**) Neutrophil numbers over the time-course of L-IVM were compared between the groups of NK46^gfp/+^ mice. Data are presented as mean fold change from the baseline (Means ± SEM, n=7-8 mice, ANOVA with Tukey’s post hoc test performed on raw data). (**C**) Representative plots of flow cytometry analysis of Ly6G surface labelling on lung NK cells (blue dots) after 4 or 24 hours after i.n. LPS application. Gated using FMO Ctrls (black dots). (**D**) Quantification per mouse of percentage of Ly6G-positive NK cells in blood, lung and spleen 4 and 24 hrs post intra nasal LPS (0.5mg/kg) compared to PBS Ctrl (n=4 mice). (**E**) Mean fluorescence intensity (MFI) of Ly6G on NK cells from mice treated with in LPS for 4 or 24 hrs (n=4 mice, Two-way ANOVA with Sidak’s post-hoc test).

Again, we could see transfer of Ly6G-positive material from neutrophils to NK cells and although imaged by L-IVM relatively rarely, it was common enough for us to see in both LPS treated and PBS control mice (Movie S4 and S5). When performing flow cytometry on the lungs of mice treated with intranasal LPS (cells stained *ex vivo* in suspension), we also found that, particularly at 4hrs after LPS, up to 25% of NK cells in the lung were positive for Ly6G compared to less than 1% in PBS treated mice (Fig. 5C and D). To our knowledge there is no expression of Ly6G mRNA in NK cells (*39*). Interestingly there was a significant decrease in Ly6G intensity on NK cells from 4hrs to 24hrs of LPS treatment, showing this transfer is markedly increased during early acute lung inflammation (Fig. 5E).

Evaluation of the same chemokines as before, at 4 hours or 24 hours post intranasal LPS application, showed equivalent levels of CXCL1 and CXCL2 in the blood and lung after 4 hrs in NK cell-depleted or IgG control mice (Fig. S4A and B). The levels of both CXCL1 and CXCL2 reverted to baseline levels more quickly in NK cell-depleted mice with levels similar to the baseline after 24 hrs (Fig. S4A and B). At later timepoints of lung inflammation, it has been reported that NK cell depletion could ameliorate the infiltration neutrophils (*40, 41*), this could be attributed to a reduction of production of IL-17 which in turn is responsible for the production of neutrophil chemo-attractants such as CXCL1 and CXCL2 (*42, 43*).

## Discussion

Tissue residency is an important feature of many cells of the immune system. There is growing appreciation that tissue resident cells, that share features and functions with circulating cells (and were at times considered developmentally-linked), can actually be phenotypically and developmentally diverse. This point is illustrated well by data on tissue resident macrophage origins (*44*). A lung resident population of NK cells has been described in humans, comprised of CD69^-^CD56^dim^ NK cells with reduced cytotoxic capabilities (*5*), a phenotype associated with more mature NK cells (*4, 45*). In the liver, a resident population of NK cells can be distinguished morphologically by electron microscopy, termed ‘pit cells’ (*46, 47*). Increased expression of the surface markers CD49a, TRAIL, CXCR6, DNAM, NKG2D, CD69 and CD200R have been used to define residency of NK cells, or ILC1, in different tissues (*48*– *50*). Alternatively, these lung intravascular ‘resident’ NK cells could be explained by an educational stage or a sentinel function of circulating NK cells (i.e. there is equilibrium with a proportion of all circulating NK cells sessile in the lung at one time). We could not find any differences in the expression of the markers above on lung NK cells compared to the blood (data not shown) consistent with the second explanation. Although leukocyte endothelium interactions have been considered to always form part of the process of diapedesis, the slow, multidirectional patrolling behaviour described for Ly6C^low^ monocytes in the vasculature of the dermis, mesentery and kidney (*51, 52*) or Ly6C^low^ monocytes and neutrophils in the glomeruli (*53, 54*) do not fit this one way progression. The intravascular NK cells did not scan the endothelium but remained stationary while continuously interacting with passing neutrophils, suggesting a different ‘leukocyte-regulatory’ function, perhaps more consistent with their innate lymphoid origin.

An explanation for the stationary NK cells would be if they were physically ‘stuck’ in the tortuous pulmonary vasculature. The alveolar capillary network has huge, highly-parallel capacity therefore, a number of capillaries ‘blocked’ in this way would likely have very little effect on the host’s capacity for gas exchange (*55*–*57*). However, neutrophils appear to be able to move freely in the same capillaries as the NK cells despite their larger diameter (12-15μm versus 10-12μm respectively). Perhaps it is actually the complex and tortuous shape of the pulmonary capillaries in combination with the high deformability of the neutrophils, particularly due to their nuclear morphology (*58*), that enables this difference in movement. Furthermore, in our intravital movies (Movies S1 and S2), clear differences in morphology between the two cell types were apparent, with NK cells being on the whole ‘round’ and neutrophils often adopting polarised or rod shapes. Neutrophil shape in LPS-stimulated mice has been associated with functional changes and neutrophil activation (*59*).

NK cells regularly probe other cells to examine their surface molecule expression and exchange of material has been described *in vitro* and indirectly *in vivo* for both human and mouse NK cells in a process termed trogocytosis (*60*–*62*). We imaged transfer of material from neutrophils to NK cells after close interaction both in L-IVM and live PCLS. Neutrophils have been described to leave ‘trails’ behind as they migrate through tissues (*63*), however, in our data all cells were intravascular. The function of Ly6G itself has not been clarified so it is unclear what role it might play in the transfer (*64, 65*). Even though it is the NK cells that acquire material from the neutrophils, trogocytotic regulation of the cells that ‘donate’ the membrane material has been demonstrated in the past, e.g. regulatory T cells ‘pruning’ costimulatory molecules from antigen presenting cells (*66*).

Our data reveal via direct analysis of cell behaviour in the lung *in vivo* that sessile, intravascular NK cells interact with neutrophils, where they can trogocytose cell surface material, influence neutrophil motility, and diminish their early accumulation in inflammation. In mice depleted of NK cells, neutrophils accumulated in greater numbers in response to an LPS ‘danger’ signal. This behaviour could not have been predicted or separated by analysing the cells in suspension, *ex vivo* or fixed tissue. It is reasonable to expect that a delicate and essential organ in close contact with the outside environment like the lung requires finessed control to prevent immunopathology in response to ‘danger’ signals in the air we breathe. Rapid neutrophil recruitment is essential for sufficient defence against many pathogens, however, superfluous inflammation can result in complications like vascular hyperpermeability. We propose that our findings signify an additional check on early neutrophil accumulation provided by sessile ‘regulatory’ NK cell behaviour during acute lung inflammation. This could therefore be potentially exploited therapeutically as a strategy to modify neutrophil behaviour in the lung without affecting the systemic neutrophil function necessary for anti-microbial defence. Our data argue that NK cells inhibit neutrophil over-accumulation in the alveolar capillaries by changing their ability to ‘scan’ the tissue, slowing them down and therefore changing the way they interact with the microenvironment.

## Materials and Methods

### Mice

NKp46^gfp/+^ transgenic mice, backcrossed to C57Bl/6J background for over 10 generations, were used (*21*). Both male and female mice between 12-20 weeks of age were used in experiments. Littermate controls were used in all experimental procedures and control and treatment mice were housed in the same cage. For L-IVM, controls and treatment mice were imaged on the same days. All mice were housed in specific pathogen free conditions at Imperial College London or the CRUK Beatson Institute, Glasgow. All experiments were carried out in accordance with the UK Animal (Scientific Procedures) Act 1986, and institutional guidelines for the care and use of animals.

### Differential leukocyte staining

NKp46^gfp/+^ mice were injected i.v. with 3μg anti-CD45 (clone 30-F11) in 100μL PBS or PBS alone. Mice were humanely killed by anaesthetic overdose (sodium-pentobarbital i.p.) 3 minutes after Ab injection and blood, lung, spleen, liver and bone marrow were harvested and processed as described below.

### Tissue preparation and flow cytometry

Blood was collected in EDTA (Thermofisher) coated syringes by cardiac puncture under terminal anaesthesia (sodium-pentobarbital i.p.). Lungs and livers were excised and then minced with scissors before incubation in DMEM (Gibco) with 10% foetal bovine serum (FBS, Gibco) at 37°C for 20 minutes with agitation before passing through a 70μm filter. Spleens were harvested and mashed through a 70μm filter. Bone marrow (BM) cells were harvested from the femur by flushing the bone with 5mL DMEM with 5% FBS and 1% Penicillin Streptomycin (Life Technologies). RBC lysis of the blood was carried out and samples were centrifuged at 450*g* for 5min at 4°C. Single cell suspensions stained Live/Dead stain (LIVE/DEAD Fixable Near-IR or Yellow, Invitrogen) and Fc-Receptor block was performed (using clone 93, BioLegend). Cell suspensions were incubated with directly conjugated fluorescent antibodies for 10 minutes at room-temperature. Count beads (CountBright, Life Technologies) were used to determine cell numbers. Compensation was performed using single stained beads (UltraComp eBeads and ArC Amine reactive compensation beads, both Thremofisher). Acquisition was performed on BDFortessa using FacsDiva software (BD Biosciences) with further analysis by FlowJo software V.10. (BD Company formerly Treestar, CA).

### Antibodies and reagents for flow cytometry

The following antibodies were used: Ly6G (clone 1A8), CD45 (clone 30-F11), CD11b (clone M1/70), CD3e (clone 145-2C11), CD19 (clone 6D5), Ter119 (clone TER-119), CD49b (clone DX5), TRAIL (clone N2B2), DNAM-1 (clone 10E5), CXCR6 (clone SA051D1), CD27 (clone LG.3A10), NKG2D (clone CX5), CD62L (clone MEL-14), CXCR2 (clone SA045E1), ICAM-1 (clone YN1/1.7.4), CD69 (clone H1.2F3), CD200R (clone OX-110) (all BioLegend) and CD49a (clone REA493, Miltenyi Biotec).

### Lung Intravital Microscopy (L-IVM)

The method used was first described by Looney et al. 2011 with a recent update (*27, 67*), modifications to this method are described below. Mice were anaesthetised with an i.p. injection of 100mg/kg Ketamine (Narketan) and 0.25mg/kg Medetomidine (Domitor) and placed on a heat mat at 37°C for the duration of the procedure. After loss-of-reflexes, Lidocaine 0.5% was injected subcutaneously at surgical sites for local anaesthesia. Mice were mechanically ventilated (MiniVent, Harvard Apparatus) with medical carbon dioxide 5% + oxygen 95%, or up to 100% oxygen, via a tracheal cannula at a 10ml/g stroke volume and 150 breaths per minute with 0.1cm/g positive end-expiratory pressure (PEEP). A custom-built flanged vacuum chamber fitted with an 8mm glass cover slip, was inserted via a 5mm long incision between 2 ribs above the left lung lobe. Minimal suction (0.1-0.2 bar) was applied to stabilise the lung against the coverslip. Imaging was performed on an upright Leica SP5 confocal microscope using a 25× 0.95na water immersion objective. Images were acquired in 3 z-slices 5µm apart with pinhole opened to give a 7.66 µm optical section (to increase light throughput and not miss cells due to the relatively low number of slices /volume needed for the imaging rate) at a frame rate of 1.6 seconds/volume. For visualization of the vasculature and different immune cell subtypes, fluorescent Abs were injected i.v. through the retro-orbital sinus whilst imaging in a maximal volume of 40μl. Anaesthesia was maintained by alternating injections of either 50mg/kg Ketamine alone or in combination with 0.125mg/kg Medetomidine at predefined timepoints. For induction of acute endotoxin induced lung injury, 1mg/kg LPS or PBS control were injected through the tracheal cannula after 20 minutes of imaging. At the end of the imaging session, mice were humanely killed by anaesthetic overdose (sodium-pentobarbital) for tissue harvest, or by cervical dislocation under anaesthesia.

### Fixed Precision Cut Lung Slices (PCLS)

NKp46^gfp/+^ mice were humanely killed by i.p. injection of sodium-pentobarbital, a small incision was made in the trachea and a customized blunted 22G needle was inserted. Subsequently, 1.2mL of 2% low-melting point agarose was instilled slowly through the needle. Mice were placed on ice for 3 min until the agarose set, and lungs were excised en-bloc and fixed in 4% formaldehyde (v/v in PBS) (VWR, UK) for 2 hrs at 4°C. Lungs were then sliced into 300μm thick sections on a vibrating microtome (Campden Ltd). Slices were stained with primary Rat anti-S100A9 (clone MU14-2A5, Hycult Biotech) and Hamster anti-CD31 Ab (2H8, Abcam UK) in PBS with 1% BSA (bovine serum albumin, Sigma), 10% NGS (normal goat serum, Sigma) and 0.001% Sodium-Azide (VWR, UK). Rat-IgG (2A3, BioXcell) and Hamster-IgG (Life Technologies) were used to control for unspecific staining. Anti-Rat Cy3 or anti-Hamster Alexa647 (Jackson) were used as secondary antibodies. Hoechst 33342 (Thermo Scientific, UK) were used to stain nuclei. Slices were imaged on a Zeiss LSM880 confocal microscope with Airyscan and further ‘Airyscan’ processed in Zeiss Software using default settings, before quantification using manual spot detection in Imaris software (Bitplane, Oxford Instuments).

### Live Precision Cut Lung Slices (PCLS)

Live PCLS procedure was equivalent to fixed PCLS with the following exceptions: 0.8mL of low-melting point agarose was used to increase the number of cells visible per FOV and limit tissue stretching. Excised lungs were places in DMEM substituted with 5% FBS and 1% Penicillin/Streptomycin (all Gibco). Lungs were immediately sliced into 300μm thick sections on a vibratome and stained with directly conjugated Ab in complete medium (phenol-red free DMEM substituted with 10% FBS, 2mM L-glutamine and 1% Penicillin/Streptomycin, all Gibco) for 30 minutes at 37°C. Slices were imaged on a Zeiss LSM880 confocal microscope with Airyscan in a full incubation chamber at 37°C with 5% CO2. The following mAb and reagents were used for live imaging: Ly6G (clone 1A8), CD31 (clone 390), CD11b (clone M1/70), CD11a (clone M17/4) (all BioLegend), Hoechst33342 (Thermo Scientific, UK). Lung slices were imaged for a maximum of 3 hrs to minimize effects from cell death and tissue damage. Airyscan time-lapse sequences processing was performed in Zeiss Software before quantification using Imaris software (Bitplane, Oxford Instuments).

### Cell tracking

L-IVM sequences were analysed using Imaris software (Bitplane, Oxford Instruments). Firstly, the full 80 minute sequence was cropped to analyse only the 20 minutes before LPS/PBS application. Subsequently an automatic gaussian filter was applied (1.01μm). NK cell tracking was performed fully manually by creating spots representing each NK cell track. For neutrophil tracking spots were initially created automatically on Ly6G-positive cells and tracks were corrected manually. XYZ position data were exported and track duration, length, speed and displacement were calculated. Cells tracked for under 3 frames (= 3.4 seconds) were excluded from the analysis. For interaction analysis NK cells and neutrophils within 15μm, measured from the cell centre, of each other for over 3.4 seconds were considered interacting. After an initial interaction a ‘buffer zone’ of 5 μm was applied, meaning the interaction was only classified as ended if the cells moved over 20 μm apart. This ‘buffer’ was applied to account for stretching/deformation of the cells during interaction, where the cell centres moved further apart, but the cells membranes were still in contact.

### Statistical and Image Analysis

Image analysis and visualization was done using Imaris (Bitplane), Fiji or Icy (Institute Pasteur). Cell tracking was performed as described above. For PCLS analysis spot detecting was performed manually and the MatLab extension ‘Spots to spots closest distance’ was used to determine the Nearest Neighbour index (NNI). Statistical analysis was performed using GraphPad Prism 7 (GraphPad Software, Inc) or in R (R Project for Statistical Computing) for the analysis of cell tracking data. Data are represented as box and whisker with all data points shown or as below. For tacking analysis, the data points represent the arithmetic means (AM) of the tracks per mouse. Results were broadly consistent regardless of whether AM, geometric means or the median were used as the measure of central tendency, and the results were not sensitive to the inclusion, omission or imputation of zero-values. The same conclusions were reached when using a generalised liner mixed effects model with nesting, a generalised linear mixed effects model without nesting, and a Box-Cox transformation followed by a linear mixed effects model. A p-value of less than 0.05 was considered significant. Details of statistical tests used can be found in the figure legends.

### NK cell depletion

To deplete NK cells, NKp46^gfp/+^ mice were injected i.p. with 200μg anti-NK1.1. (Clone PK136, BioXcell, NH, USA) or IgG Control (Clone C1.18.4, BioXcell, NH, USA) in 100ul PBS at 48 hours and 2 hours before LPS/PBS administration. Efficiency of the depletion was assessed by flow cytometry.

### LPS-induced acute lung inflammation without L-IVM

NKp46^gfp/+^ mice were anaesthetized by isoflurane inhalation and 0.5mg/kg LPS or PBS control at a total volume of 50μl were administered intranasally. Mice were left to recover fully from anaesthesia and were monitored at regular intervals. At indicated timepoints (4h – 24h) mice were humanely killed by anaesthetic overdose (sodium-pentobarbital) for blood and tissue harvest.

### Quantification of cytokine expression

Enzyme linked immunoassay of CXCL1 and CXLC2 (both R&D Systems) were performed on plasma, lung and spleen supernatants on NKp46^gfp/+^ mice treated with anti-NK1.1 or IgG Control as described above. All procedures were performed according to the manufacturer’s instructions. Luminescence measures were performed on a Tecan Sunrise analyser.

## Acknowledgments

### General

We thank Gerry Graham, Cristina Lo Celso, Martin Spitaler, Sarah Rankin and Clare Lloyd for expertise, help and useful discussion of the work. Microscopy and cell tracking were performed in the Beatson Advanced Imaging Resource and Imperial College Facility for Imaging by Light Microscopy (in part funded by Wellcome Trust grant 104931/Z/14/Z). We thank Catherine Winchester and Laura Machesky for help with editing the manuscript.

### Funding

J.S. is supported by an Imperial College London President’s PhD Scholarship. K.D.F. is funded by Wellcome Trust (201356/Z/16/Z). J.B.G.M is supported by the NHLI foundation PhD studentship, Imperial College London. J.V. is funded by Emil Aaltonen Foundation, Sigrid Juselius Foundation and Jane and Aatos Erkko Foundation. X.L.R. and A.J.M. are core funded by Cancer Research UK (A23983 and A17196). N.G. is supported by the Wellcome Trust (RCDF088381/Z/09/Z). L.M.C. is funded by core support from Cancer Research UK (A23983 and A17196), the MRC (MR/M01245X/1), and National Heart & Lung Institute Foundation M.N. is funded by core support from Cancer Research UK.

### Author contributions

J.S. designed and performed the experiments, performed analysis and wrote the manuscript. K.D.F., J.B.G.M., J.V., X.L.R. and A.J.M. helped with experiments and contributed to interpretation. K.D.F. performed cytokine assays. M.N. performed statistical analysis and mathematical modelling. N.G., M.B.H., M.F.K. helped with interpretation, provided expertise, reagents and edited the manuscript. L.M.C. conceptualised and directed the project, obtained funding, designed experiments, contributed to interpretation and wrote the manuscript. All authors reviewed and approved the manuscript.

### Competing interests

The authors declare that they have no competing interests.

### Data and materials availability

Request for access to this data can be made to the corresponding author. All other data is provided in the paper or in the Supplementary Materials.

