## Supplemental Material for "Pulmonary natural killer cells control neutrophil intravascular motility and response to acute inflammation"

**Affiliations:**

[6]Current address, Fred Hutchinson Cancer Research Centre, Seattle, WA, USA.

**This PDF file includes:**

Figs. S1 to S4
Tables S1 to S3
Captions for movies S1 to S6

**Other supplementary materials for this manuscript include the following:**

Movies S1 to S6

### Supplementary Figures

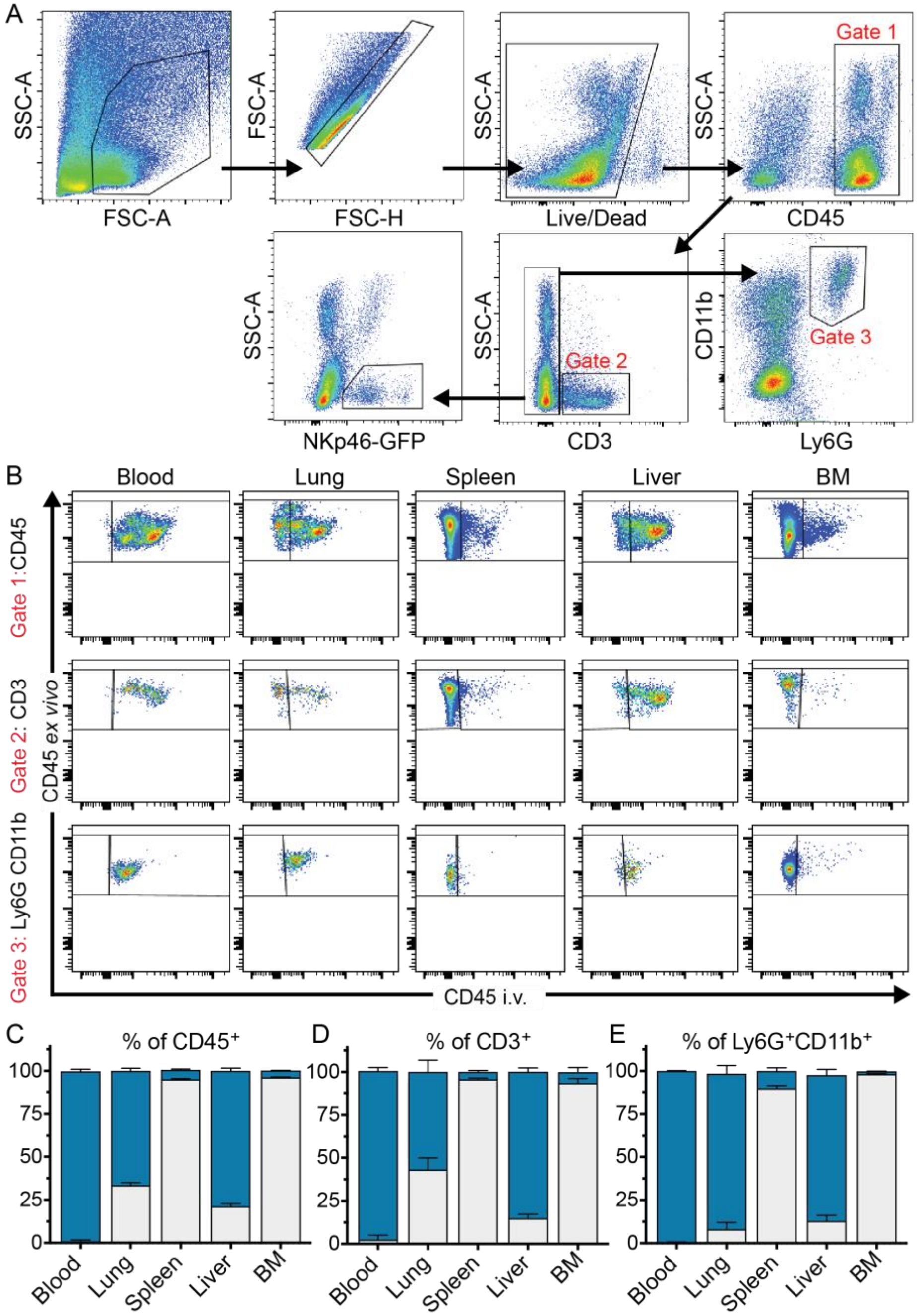


**Fig. S1.** Flow cytometry gating strategy for NK cells and neutrophils from murine blood. (**A**) Cells were selected for singlets and gated on Zombie Yellow negative, CD45 positive population. 1A8 clone against Ly6G and clone M1/70 against CD11b were used to identify neutrophils. NK cells were defined by GFP expression and lack of CD3. Analysis of vascular versus tissue distribution of different leukocyte subsets (SSC = side scatter; FSC = forward scatter; A = area; H = height). (**B**) Representative flow cytometry plots of total CD45+ cells, T cells (CD3+) and neutrophils (Ly6G+, CD11b+) in blood, lung, spleen, liver and BM of NKp46$^{gfp/+}$ mice injected with 3ug anti-CD45 Alexa700 AB. (**C-E)** Quantification of flow cytometry data showing the percentage of intravascular cells (blue) and parenchymal cell (grey). (n=3, error bars represent SD)

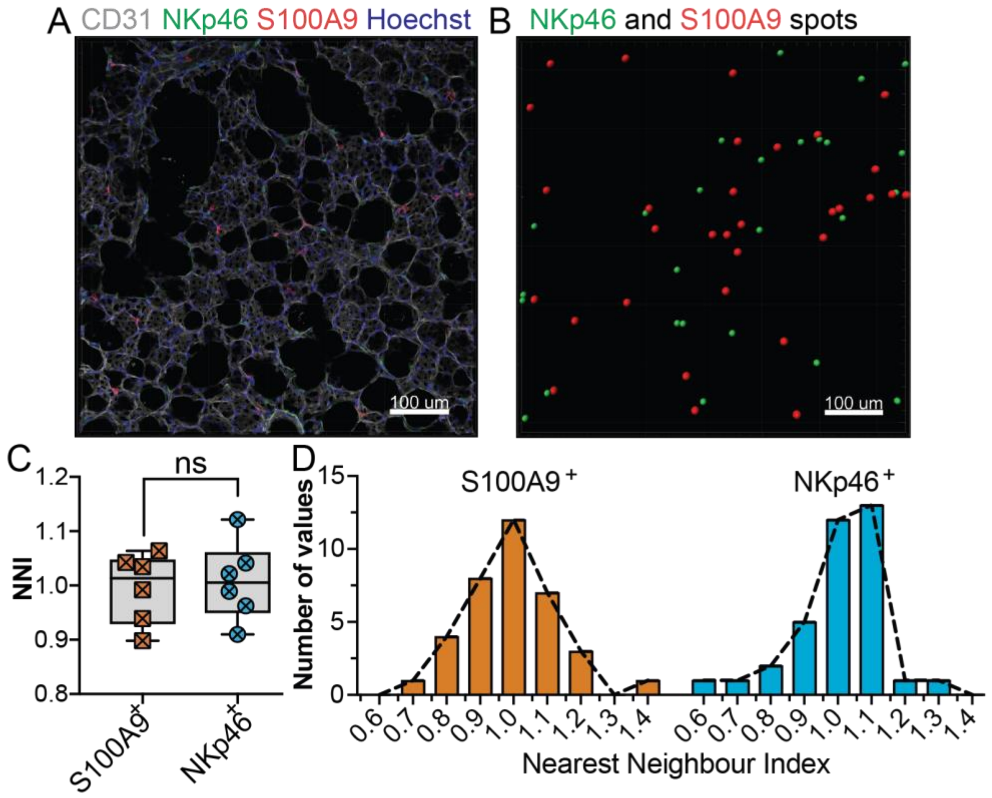


**Fig. S2.** Distribution of NK cells and neutrophils in the lung. (**A**) Representative images of lung PLCS from Ncr1$^{gfp/+}$ mice. Fixed slices were stained with anti-CD31 (gray), anti-S100A9 (red) and Hoechst (blue), to label the vasculature, neutrophils and nuclei respectively. (**B**) Imairs (Bitplane) was used to segment NK cells (green) and S100A9-positive neutrophils (red). (**C**) Nearest-neighbour index (NNI) was used to determine the spatial distribution of neutrophils and NK cells in the lung parenchyma. (**D**) Histograms of the cell distribution (n=6 mice with 6 images per lung, Unpaired Student's t-test)

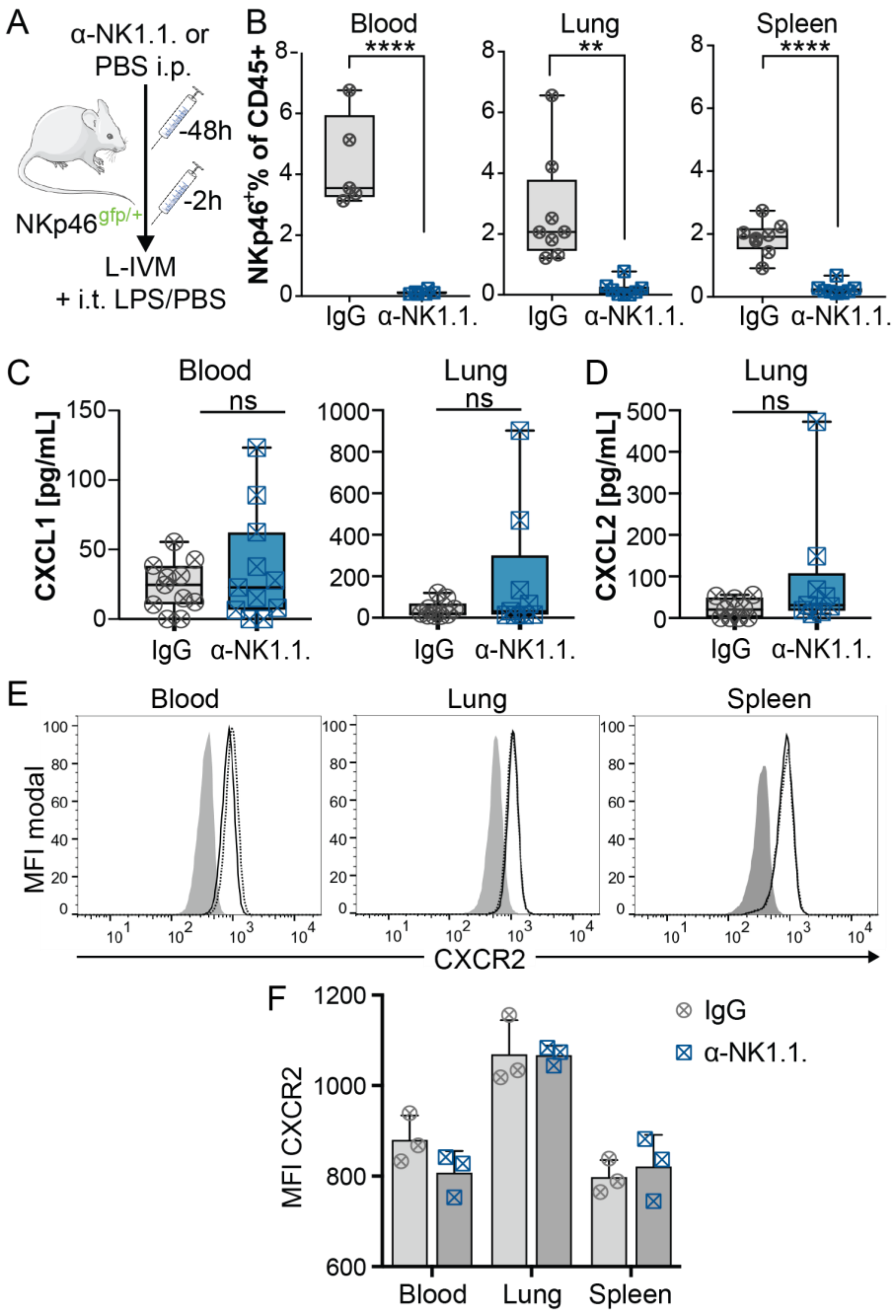


**Fig. S3.** Antibody mediated NK cell depletion does not alter neutrophil chemokine parameters in a steady-state. (**A**) Timeline of anti-NK1.1. or IgG injections for NK cell depletion. (**B**) Depletion efficiency was confirmed by flow cytometry in independent experiments (n=5 mice for blood and 8 mice for lungs and spleen, Unpaired two-tailed t-tests; $**P < 0.005$, $****P < 0.00005$). Analysis of CXCL1, CXCL2 and CXCR2 expression after NK cell depletion. CXCL1 (**C**) and CXCL2 (**D**) levels were quantified by ELISA of whole blood and lung supernatants (n=11 mice per condition from 4 independent experiments; Unpaired two-tailed t-tests with Welch's correction). For CXCR2 expression analysis neutrophils were gated based on expression of Ly6G and CD11b. (**E**) Representative histograms of CXCR2 expression on neutrophils from anti-NK1.1. treated mice (black line) or IgG Ctrls (dotted line) compared to FMO controls (grey). (**F**) Quantification of CXCR2 expression on neutrophils from anti-NK1.1. or IgG Ctrls treated mice (n=3).

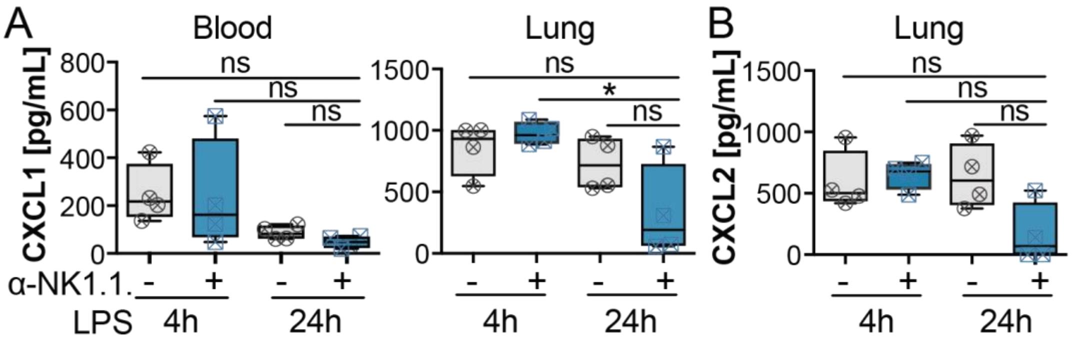


**Fig. S4.** ELISA were performed to quantify cytokines after endotoxin treatment. Levels of CXCL1 (**A**) and CXCL2 (**B**) in whole blood or lung supernatants of mice with intranasal application of LPS for 4 hrs or 24hrs (n=4 mice; ANOVA with Tukey's post hoc test; * $P < 0.05$).

**Table. S1.** Paired t-test comparing NK-interacting, with non-interacting neutrophil tracks in IgG treated mice.

| ***Measure*** | ***Unit*** | ***Estimated difference*** ***Interacting – non-interacting*** | ***95% CI Lower*** | ***95% CI Upper*** | ***p-value*** |
|---|---|---|---|---|---|
| ***Duration*** | [min] | 4min 35sec | 3min 40sec | 5min 30sec | 0.0001 |
| ***Length*** | [µm] | 133.483 | 85.813 | 181.153 | 0.0008 |
| ***Speed*** | [µm/s] | -1.806 | -3.078 | -0.534 | 0.0148 |
| ***Displacement*** | [µm] | 13.199 | 3.648 | 20.750 | 0.0163 |

**Table S2.** Comparison of neutrophil track parameters around the interaction with an NK cell

| ***Measure*** | ***Unit*** | ***Comparison*** | ***Estimated difference*** | ***95% CI Lower*** | ***95% CI Upper*** | ***p-value*** |
|---|---|---|---|---|---|---|
| ***Duration*** | [min] | During - Before | -0min 51sec | -2min 15sec | 0min 33sec | 0.1774 |
| | | After - During | 0min 24sec | -0min 54sec | 1min 43sec | 0.4667 |
| ***Length*** | [µm] | During - Before | -28.521 | -77.373 | 20.330 | 0.1937 |
| | | After - During | 23.963 | -18.644 | 66.571 | 0.2079 |
| ***Speed*** | [µm/s] | During - Before | 0.079 | -0.139 | 0.297 | 0.3938 |
| | | After - During | -0.0147 | -0.233 | 0.203 | 0.8688 |
| ***Displacement*** | [µm] | During - Before | -7.294 | -15.605 | 1.017 | 0.0737 |
| | | After - During | 6.738 | -1.784 | 15.261 | 0.0978 |

**Table. S3.** Welch's t-test of IgG treated, non-interacting neutrophils compared to neutrophils in NK cell-depleted mice.

| *Measure* | *Unit* | *Estimated difference IgG – α-NK1.1.* | *95% CI Lower* | *95% CI Upper* | *p-value* |
|---|---|---|---|---|---|
| ***Duration*** | [min] | 0min 14sec | -1min 26sec | 1min 54sec | 0.7537 |
| ***Length*** | [µm] | -10.118 | -58.297 | 38.061 | 0.6499 |
| ***Speed*** | [µm/s] | -1.173 | -3.788 | 1.442 | 0.3326 |
| ***Displacement*** | [µm] | -2.561 | -14.448 | 9.325 | 0.6320 |

**Movie S1. NK cells remain stationary within pulmonary vessels for long periods.** L-IVM of NKp46$^{gfp/+}$ mouse. NK cells (green) express GFP, whereas neutrophils (orange) and vasculature (gray) are visualized by iv injection of fluorescently labelled anti-Ly6G (1A8) and Isolectin-B4 respectively. Movement at 20 minutes due to intratracheal instillation of LPS.

**Movie S2. Neutrophil behavior in the lung in a steady state.** L-IVM of C57Bl/6 mice injected iv with fluorescently labelled anti-Ly6G (1A8) and Isolectin-B4, to visualize neutrophils (red) and vasculature (gray) respectively. Intratracheal instillation of PBS after 30 minutes of baseline imaging.

**Movie S3. Material is transferred from a neutrophil to an NK cells after an interaction.** L-IVM of NKp46$^{gfp/+}$ mouse. NK cells (green) express GFP, whereas neutrophils (orange) and vasculature (gray) are visualized by iv injection of fluorescently labelled 1A8 and Isolectin-B4 respectively.

**Movie S4. Transfer occurs in mice treated with murine IgG antibody.** L-IVM of NKp46$^{gfp/+}$ mouse. NK cells (green) express GFP, whereas neutrophils (orange) and vasculature (grey) are visualized by iv injection of fluorescently labelled 1A8 and Isolectin-B4 respectively. Intratracheal instillation of PBS after 20 minutes of baseline imaging.

**Movie S5. Transfer occurs more after acute LPS instillation.** L-IVM of NKp46$^{gfp/+}$ mouse. NK cells (green) express GFP, whereas neutrophils (orange) and vasculature (gray) are visualized by i.v. injection of fluorescently labelled 1A8 and Isolectin-B4 respectively. Intratracheal instillation of LPS after 20 minutes of baseline imaging.

**Movie S6. Neutrophils accumulate rapidly in the lung after LPS instillation.** L-IVM of C57Bl/6 mice injected i.v. with fluorescently labelled anti-Ly6G (1A8) and Isolectin-B4, to visualize neutrophils (red) and vasculature (gray) respectively. Intratracheal instillation of LPS after 30 minutes of baseline imaging.
